# Determinants of DDX3X sensitivity uncovered using a helicase activity in translation reporter

**DOI:** 10.1101/2023.09.14.557805

**Authors:** Kevin C. Wilkins, Till Schroeder, Sohyun Gu, Jezrael L. Revalde, Stephen N. Floor

**Affiliations:** Department of Cell and Tissue Biology, University of California, San Francisco, San Francisco, California, 94143, USA; Graduate Division, University of California, San Francisco, San Francisco, CA, United States; Julius-Maximilians-University of Würzburg, Würzburg, 97070, Germany; Department of Pharmaceutical Chemistry, University of California, 600 16th Street, San Francisco, California 94143, United States; Helen Diller Family Comprehensive Cancer Center, University of California, San Francisco, San Francisco, California, 94143, USA

**Keywords:** RNA Helicase, DDX3X, Ribosome scanning, DEAD-box helicase, Translation initiation, mRNA

## Abstract

DDX3X regulates the translation of a subset of human transcripts containing complex 5′ untranslated regions (5′ UTRs). In this study we developed the helicase activity reporter for translation (HART) which uses DDX3X-sensitive 5′ UTRs to measure DDX3X mediated translational activity in cells. To dissect the structural underpinnings of DDX3X dependent translation, we first used SHAPE-MaP to determine the secondary structures present in DDX3X-sensitive 5′ UTRs and then employed HART to investigate how their perturbation impacts DDX3X-sensitivity. Additionally, we identified residues 38-44 as potential mediators of DDX3X’s interaction with the translational machinery. HART revealed that both DDX3X’s association with the ribosome complex as well as its helicase activity are required for its function in promoting the translation of DDX3X-sensitive 5′ UTRs. These findings suggest DDX3X plays a crucial role regulating translation through its interaction with the translational machinery during ribosome scanning, and establish the HART reporter as a robust, lentivirally encoded measurement of DDX3X-dependent translation in cells.

## INTRODUCTION

DDX3X is a ubiquitously expressed RNA-helicase implicated in almost all stages of RNA metabolism^1,2^. By influencing gene expression, RNA localization, and RNA stability, DDX3X impacts cellular functions and responses, including homeostasis^3^, stress granule regulation^4,5^, cell stress^6,7^, signaling^1^, the immune response^8,9^, and embryonic development^10,11^. Dysfunctions in DDX3X are linked with several human diseases, including cancer, viral infection, inflammation, and intellectual disabilities^1,12^.

DDX3X is a member of the DEAD-box RNA helicase family and hydrolyzes ATP to unwind RNA molecules and remodel RNA-protein complexes^13^. DDX3X shows high conservation with its yeast homolog Ded1,with whom it forms a clearly defined subfamily^14^. DDX3X/Ded1 helicases are characterized by a conserved helicase core domain which contains nine motifs involved in ATP binding, hydrolysis, and RNA interaction^1,13^. While the helicase core is highly conserved across the DEAD-box family, the N- and C-termini are variable^13,15^. These termini are involved in protein-protein interactions and subcellular localization, but their role is yet to be fully understood^1^.

DDX3X plays a role in translation by unwinding RNA secondary structures and remodeling RNA-protein complexes. DDX3X/Ded1 have been suggested to interact with components of the translation initiation machinery, including the eIF4F complex (eIF4E, eIF4G, and eIF4A), to promote ribosome binding to the mRNA’s 5’ cap and enhance the recruitment of ribosomes to the mRNA^1,4,16–18^. During ribosome scanning, DDX3X unwinds secondary structures within the 5’ UTRs to facilitate the movement of ribosomes along the mRNA to allow for identification of the start codon^2,19^. One regulatory feature DDX3X might modulate is the expression of uORF, since Ded1 has been shown to regulate translation of near cognate uORF^16^. As opposed to eIF4A, a generalist necessary for global translation, DDX3X has been observed to act as specialist helicase, required for the proper translation of a subset of the human mRNAs^13,19^. This specialized role could derive from DDX3X/Ded1’s higher RNA unwinding capability compared to eIF4A, suggesting that DDX3X might be required to resolve complex and highly structured 5’ UTRs^20^.

Several aspects of DDX3X’s activity during ribosome scanning and translation are still poorly understood. The complete repertoire of its target mRNAs and their diversity across cell types and contexts is not fully characterized, nor is it understood what 5’ UTRs features and structures confer dependency on DDX3X for proper translation regulation. Further, while DDX3X has been observed to interact with helix 16 of the 18S rRNA, a component of the small ribosomal subunit^19^ the function and extent of its association with the ribosome and other translation factors is still unclear. Given the substrate-dependency of DDX3X’s role in diseases, a better picture of such variables is necessary for insights into the broader landscape of translation regulation and its implications in health and disease.

In this paper, we created a helicase activity reporter for translation (HART) to measure the translational activity of DDX3X. We characterized the structures and features of 5’ UTRs who rely on DDX3X for proper translation and employed HART to dissect this sensitivity to DDX3X. We then identified residues in the N-terminus of DDX3X as mediating interaction with the ribosomal complex and employed HART to determine that this interaction is important for DDX3X’s role in promoting translation. Together, our work constitutes progress in the understanding of the mechanism of DDX3X-mediated translation and the creation of a versatile reporter with potential implications in both further dissection of DDX3X function and drug discovery.

## RESULTS

### Creating a reporter for the translational activity of DDX3X

We set out to create a reporter to investigate the translational activity of DDX3X. We first identified human transcripts whose translation is dependent on DDX3X. Ribosome profiling in prior work showed that the loss of DDX3X caused a decrease in ribosome density on a subset of transcripts, including ODC1, RAC1, and HMBS^19^. The 5′ untranslated regions (5′ UTR) of these transcripts requires DDX3X for efficient translation in in vitro and in cell assays^19,21^. We define these sequences as DDX3X-sensitive 5′ UTRs and reasoned we could use them as the foundation of a reporter for DDX3X translation activity in cells (Figure 1A, B).

**Figure 1:**
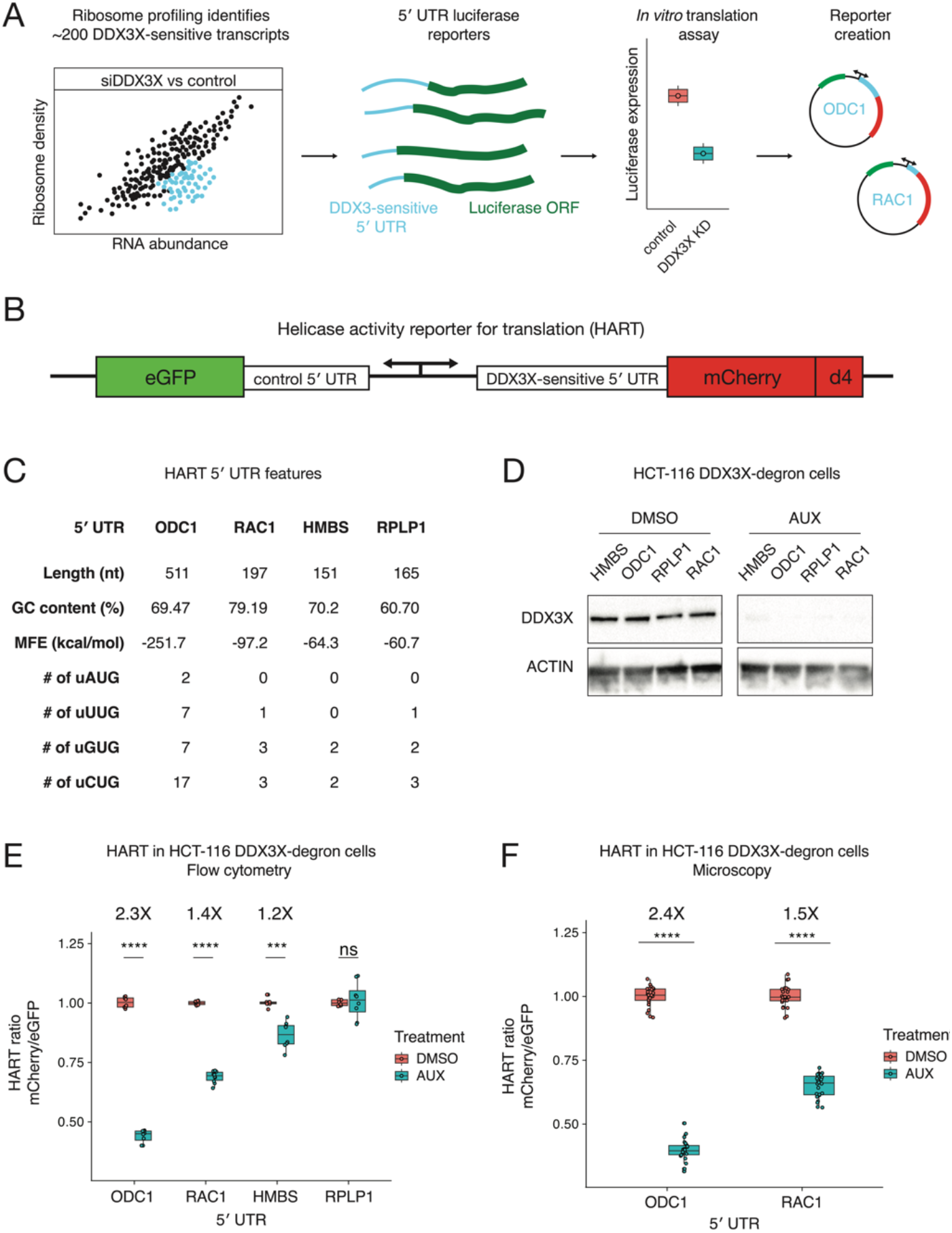
The helicase activity reporter for translation (HART) employs DDX3X-sensitive 5′ UTRs to measure the translational activity of DDX3X. (**A)** Selection of DDX3X-sensitive 5′ UTRs for HART. Validated DDX3X-sensitive 5′ UTR from prior ribosome profiling and in vitro translation, as well as negative controls, were cloned into the HART reporter. **(B)** Diagram of HART. HART is constructed around a bidirectional promoter, with one arm directing the transcription of a control 5′ UTR and eGFP, and the other arm featuring a DDX3X-sensitive 5′ UTR followed by mCherry. An ornithine decarboxylase (ODC) degron (d4) shortens the mCherry half-life. The HART ratio (mCherry/eGFP) can be used to measure the translational activity of DDX3X. **(C)** Table of features for the 5′ UTRs from C. Features catalogued include length, GC content, RNA minimum free energy (MFE) prediction using the ViennaRNA Package^48^, and the number of upstream AUGs and near cognate codons. **(D)** Western blot for cells in A. HCT116 degron cells were treated with either auxin or DMSO, and then lysates were collected and resolved by SDS-page and used for western blotting with antibodies against actin and DDX3X. **(E)** HART ratio in HCT116 degron cells analyzed with flow cytometry. HCT116 degron cells were lentivirally transduced with HART constructs with indicated 5′ UTRs upstream of mCherry. After 48 hours from the addition of either DMSO or auxin, which induces degradation of endogenous DDX3X, the fluorescent signal of cells was measured by fluorescent cytometry. The HART ratio (mCherry/eGFP) was calculated for each cell and averaged across replicate wells. Statistical significance was determined by unpaired t-test: ns: p > 0.05, *: p <= 0.05, **: p <= 0.01, ***: p <= 0.001, ****: p <= 0.0001. **(F)** HART ratio in HCT116 degron cells analyzed by microscopy. HCT116 degron cells were transduced with HART construct for ODC1 or RAC1. After 48 hours from the addition of either DMSO or auxin the cells were fixed and imaged using an InCell Analyzer 6500HS. The HART ratio was calculated for each cell and averaged across replicate wells. Statistical significance was determined by unpaired t-test: ns: p > 0.05, *: p <= 0.05, **: p <= 0.01, ***: p <= 0.001, ****: p <= 0.0001.

We selected four 5′ UTRs, ODC1, RAC1, HMBS, and RPLP1 possessing differing features (Figure 1C) and differing translational sensitivity to DDX3X^19^. We incorporated these 5′ UTR into a construct we called helicase activity reporter for translation (HART, Figure 1B). HART is a lentiviral plasmid featuring a dual promoter, composed of a copy of human PGK (hPGK) and one of mini CMV, which efficiently directs translation in a bidirectional fashion both in cells and *in vivo*^22^. We used hPGK to direct transcription of a DDX3X-sensitive 5′ UTR followed by the open reading frame (ORF) of the fluorescent protein mCherry. The mini CMV promoter was used to direct transcription of a DDX3X-insensitive 5′ UTR followed by the ORF of eGFP to act as an internal control. The DDX3X-insensitive 5′ UTR consists of a short 9 nucleotides (nt) sequence (CTAGCCACC) which was previously used as a control in cellular and in vitro assays of DDX3X-sensitive translation^19^. In order to better detect changes in protein levels, we fused mCherry with the ornithine decarboxylase (ODC) degron (d4) known to confer a protein half-life of 4 hours^23^. In the HART reporter, mCherry protein levels are a proxy for translation of DDX3X-sensitive transcripts, while eGFP protein levels are independent of DDX3X changes and act as an internal control. The mCherry/eGFP ratio, which we refer to as the HART ratio, is thus a readout of the ability of DDX3X to promote translation.

To validate HART as a reporter, we assessed the impact of DDX3X loss on the HART ratio. We depleted endogenous DDX3X using an inducible degron system, employing a male-derived colorectal cancer HCT 116 cell line with stably integrated auxin-activated OSTIR1 machinery, where the endogenous DDX3X gene was tagged with a degradation tag^19,24^. Treating the cells with auxin for 48 hours resulted in near complete degradation of endogenous DDX3X protein (Figure 1D). We transduced HART into these degron cells and treated them with auxin for 48 hours. Flow cytometry measurements of the ratio of mCherry to eGFP signal identified a significant decrease in auxin-treated cells compared to the DMSO-treated control in HART constructs containing HMBS, ODC1, or RAC1 but not RPLP1 5′ UTRs (Figure 1E, Supplementary Figure 1A). These results were in line with previous in cell and in vitro translation assays^19,25–27^. Notably, there was no difference in eGFP fluorescence levels between auxin-treated cells and controls, while mCherry levels exhibited significant differences, demonstrating that the change in HART ratio between treatments was driven by the DDX3X-sensitive 5′ UTR (Supplementary Figures 1B-C). Microscopy measurements were consistent with flow cytometry, confirming the reliability and generalizability of HART-ODC1 and HART-RAC1 (Figure 1F). Overall, these results collectively demonstrated the effectiveness of HART as a reporter for evaluating DDX3X’s impact on translation initiation.

### Structural probing of the RAC1 and ODC1 5′ UTRs

DDX3X is necessary for the proper translation of several human mRNAs, but the precise understanding of which 5′ UTR structures and features render a given transcript sensitive to DDX3X remains incomplete. Prior work has implicated upstream start codons, RNA structure, GC content, and other mRNA features as important in DDX3X-controlled regulation^19,25,28,29^. However, experimental evidence of RNA structure in DDX3X-sensitive mRNAs is limited. To gain deeper insights into the elements that confer DDX3X sensitivity to the RAC1 and ODC1 5′ UTRs, we used SHAPE-MaP structure probing to experimentally determine their secondary RNA structures^30,31^.

To measure RNA structures using SHAPE-MaP, mRNAs containing the RAC1 and ODC1 5′ UTRs and the luciferase ORF were in vitro transcribed using T7 polymerase^19,21^. RNA was then folded, probed with NAI or DMSO as control, and then reverse transcribed and sequenced. Mutation profiles and SHAPE reactivity were measured and used to model RNA secondary structures (Figure 2 A-D).

**Figure 2:**
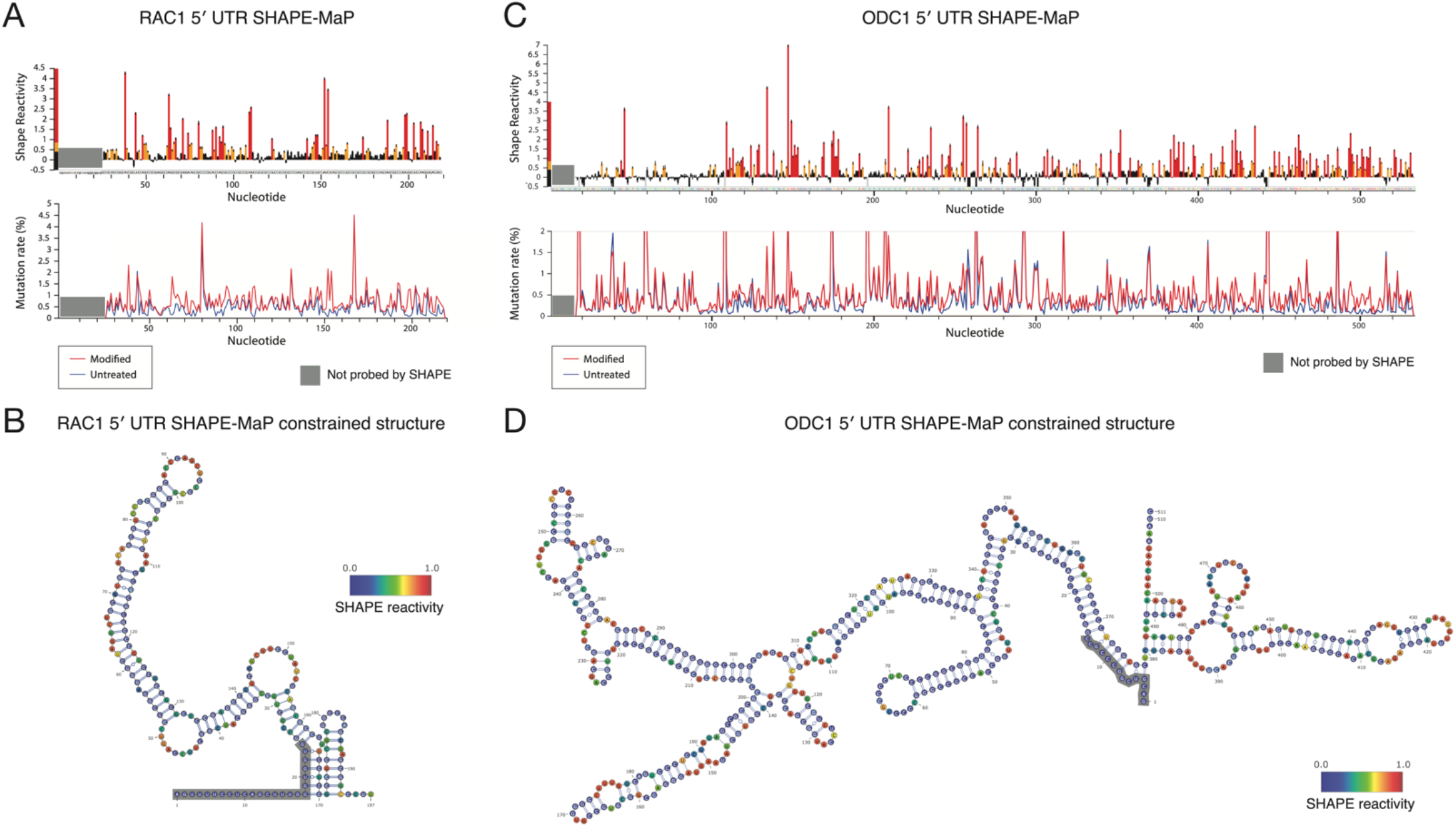
DDX3X sensitive 5′ UTRs are highly structured. **(A)** SHAPE-MaP reactivity and mutation rate for the 5′ UTR of RAC1 *in vitro. In vitro* transcribed mRNA containing the 5′ UTR of RAC1 and the open reading frame of luciferase was probed with 200 mM NAI or DMSO control for SHAPE-MaP. The RNA was reverse transcribed, sequenced, and analyzed to obtain mutation profiles and SHAPE reactivity with the ShapeMapper tool. **(B)** Diagram of the structure of the RAC1 5′ UTR *in vitro*, based on data from Figure 2A and computed with ShapeMapper 2.1.3. (Busan & Weeks, 2018) **(C)** SHAPE-MaP reactivity and mutation rate for the 5′ UTR of RAC1 *in vitro. In vitro* transcribed mRNA containing the 5′ UTR of ODC1 and the open reading frame of luciferase was probed with 200 mM NAI or DMSO control for SHAPE-MaP. The RNA was reverse transcribed and sequenced. The mutation rate was calculated at each position for both treated and control samples. The SHAPE reactivity was calculated based on the difference in mutation rate. SHAPE reactivity is cropped for space, the full figure can be found in Supplementary Figure 2A. **(D)** Diagram of the structure of the ODC1 5′ UTR *in vitro*, based on data from Figure 2C and computed with ShapeMapper 2.1.3. (Busan & Weeks, 2018)

RNA structure prediction constrained by the SHAPE-MaP data for the RAC1 5′ UTRs predicts a complex structure, comprising a 152 nt long complex structure and a 23 nt stem loop (Figure 2A-B). The RAC1 5′ UTR has a 79% GC content and a predicted free energy of -97.20 kcal/mol. The RAC1 SHAPE-MaP result obtained in vitro is alike the one obtained in cells (Supplementary Figure 2 B-C) or via in silico predictions using the RNAfold software (Supplementary Figure 2D). The ODC1 5′ UTR also showed a highly complex structure, with a long 376 nt structure with several smaller hairpins branching from it and a shorter 107 nt structure, with an overall free energy of - 251.70 kcal/mol (Figure 2C-D, Supplementary Figure 2A) and a 69% GC content. These in vitro probing results are overall similar to RNAfold prediction (Supplementary Figure 2E).

The presence of large stem loops and overall high complexity of these 5′ UTRs could impede efficient translation initiation in the absence of the unwinding and remodeling activity of helicases^32^. DDX3X may be crucial in resolving these stable stem loops and enabling mRNA accommodating into the small ribosomal subunit or scanning across the 5′ UTR to identify the translation start site. Previous work has suggested that Ded1, the yeast homolog, has a higher unwinding activity than eIF4A, which may position Ded1/DDX3X as a specialist helicase required to unwind particularly complex structures^20^.

### Dissection of RAC1 and ODC1 DDX3X-sensitivity

We next sought to determine which features of the ODC1 and RAC1 5′ UTRs contribute to DDX3X-sensitivity. To explore this in an unbiased manner, we created HART constructs with five equal size deletions tiling the 5′ UTRs of both RAC1 (Figure 3A and B) and ODC1 (Figure 3A and E). These constructs were then used to measure the HART ratio in HCT116 degron cells after 48 hours of auxin or DMSO treatment.

**Figure 3:**
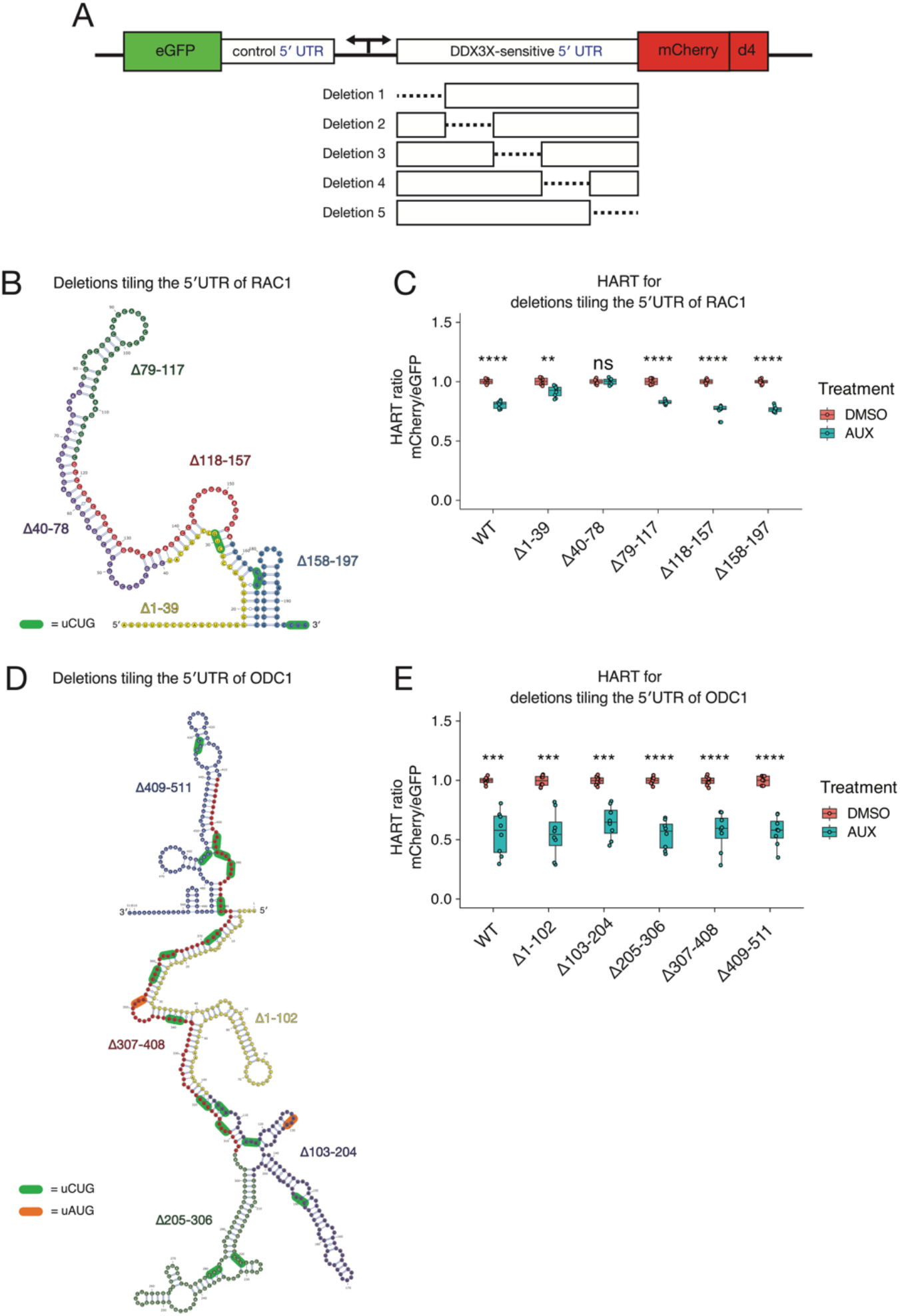
Dissection of RAC1 and ODC1 DDX3X-sensitivity. **(A)** Schematic of the HART construct with deletions tiling DDX3X-sensitive 5′ UTRs. **(B)** Diagram of the structure of RAC1 with the deletions highlighted. **(C)** HART data for panel B. HART constructs containing RAC1 5′ UTR or five deletion mutants were transduced via lentivirus into HCT116 degron cells. The cells were treated with auxin to induce loss of endogenous DDX3X or DMSO control for 48 hours, and the HART ratio (mCherry/eGFP) was measured by flow cytometry. **(D)** Diagram of the structure of ODC1 with the deletions highlighted. **(E)** Same experiment as in C, but with HART-ODC1 constructs from D.

Deletions of the 79-117, 118-157, and 158-197 regions of the RAC1 5′ UTRs residues conferred similar DDX3X-sensitivity compared to full length wildtype. However, deletion of the 1-39 bases led to diminished sensitivity, while deletion of the 40-78 bases showed an even further reduction to the point where there was no statistical difference between cells treated with auxin or DMSO. The 1-39 and 40-78 deletions impact the large structure in the RAC1 5′ UTR, suggesting that unwinding of this structure by DDX3X is important in the scanning of its 5′ UTRs (Figure 3B). This is consistent with prior work that identified an important role for cap-proximal RNA structures in DDX3X-mediated translation^29^. Notably, in silico folding predicts the Δ40-78 RAC1 5′ UTRs possess the lowest free energy among all the deletions, suggesting that a less stable structure exhibits decreased sensitivity to DDX3X (Supplementary Figure 3). In contrast, we find that deletions in the ODC1 5′ UTR do not impact DDX3X sensitivity, suggesting its overall GC content or structure may be sufficiently complex that no single deletion ablates a requirement for DDX3X (Figure 3G). Taken together, we find that DDX3X sensitivity is mediated by regions within some but not all DDX3X-sensitive mRNAs.

### The 38-44 residues of DDX3X contribute to its association with the translational machinery

The dynamics of DDX3X unwinding of 5′ UTR structures during ribosome accommodation and/or scanning are currently unknown. One possibility is that DDX3X unwinds structures ahead of ribosome scanning, independently and untethered from the ribosome or other ribosome-associated factors in a trans fashion. Alternatively, it is possible that DDX3X is bound or tethered to the ribosome during ribosome scanning and that it unwinds 5′ UTRs while tethered, in a cis fashion. The cis scenario would be alike the behavior of related DEAD-box RNA helicase eIF4A which associates with the ribosome during scanning^33^. Some evidence for the cis model is provided by iCLIP data, which suggests that DDX3X binds to the helix 16 of the 18S rRNA in the small subunit of the ribosome, which is located near the mRNA entry channel^19^. Interestingly, a similar result was also found in yeast, where iCLIP identified helix 16 as crosslinking with Ded1, the yeast homolog of DDX3X^16^.

While helix 16 is a potential location of binding of DDX3X to the ribosome, it is unknown what part of the DDX3X protein would be responsible for this interaction. DDX3X is constituted by two RecA-like domains (domain I and domain II) that make up the helicase core, which is highly conserved across the DEAD-box helicase family and across species. The helicase core is flanked by the N-terminal extension (NTE) and the C-terminal extension (CTE), which were found to also be essential to the protein’s functional helicase activity^15,34^. Finally, the N-terminus and the C-terminus are present at the extremities of the protein and are significantly less conserved across the protein family and across species (Figure 4A), although contain several regions conserved across the DDX3X/Ded1 subfamily^1^.

**Figure 4:**
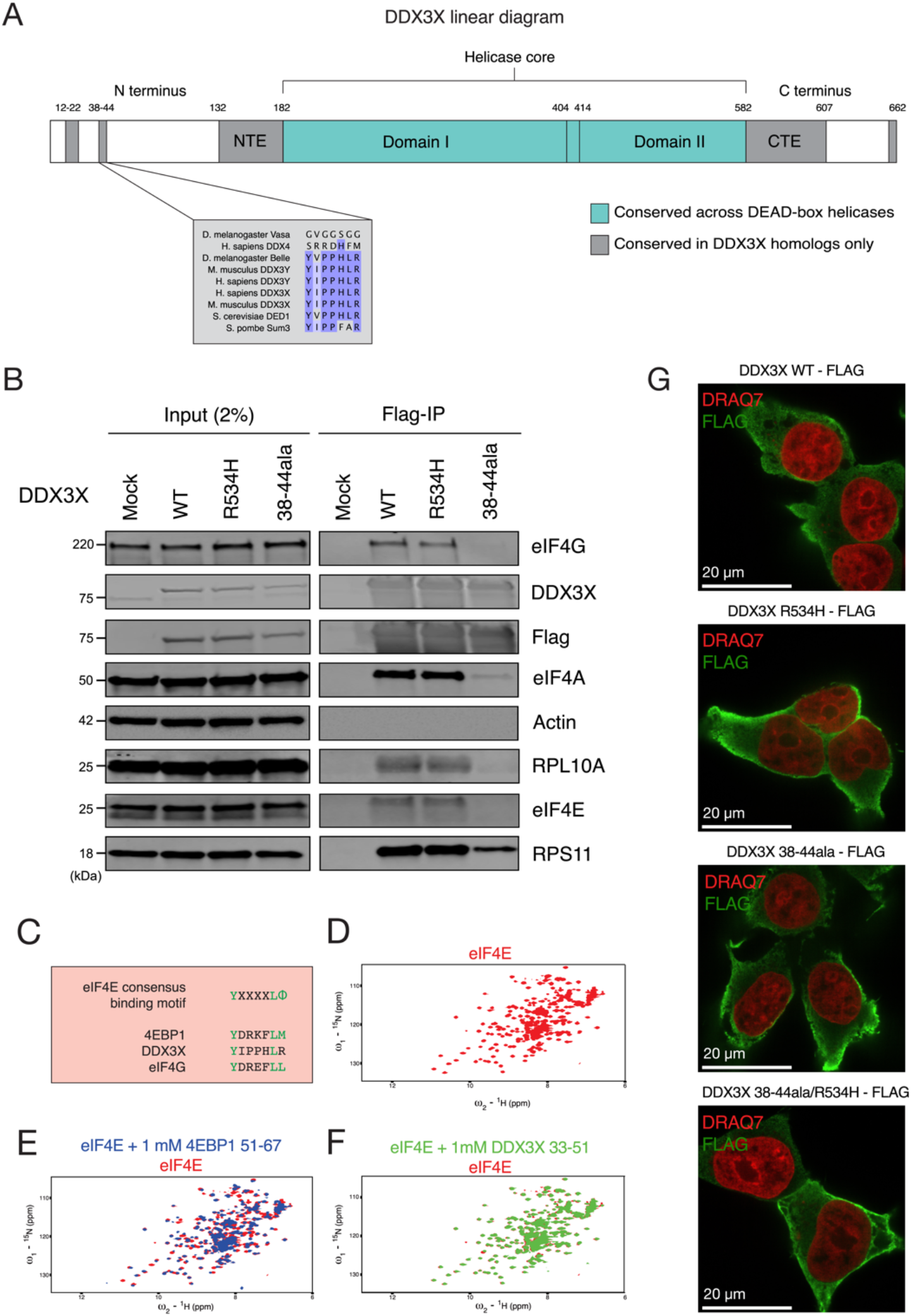
DDX3X interacts with the translational machinery via the 38-44 residues. **(A)** Diagram of the DDX3X protein structure. DDX3X contains a helicase core, highly conserved across the DEAD-box helicase protein family, composed of two RecA-like domains, denoted as Domain I and Domain II (denoted in blue). In addition to the helicase core, the functional core of the protein also includes the NTE and CTE, which have been shown to be necessary for the RNA unwinding activity of DDX3X. Outside of the functional core there are the N- and C-termini, which are less conserved across the protein family, but contain regions conserved across the Ded1/DDX3X subfamily (denoted in gray). **(B)** DDX3X co-immunoprecipitation with translation machinery proteins. HEK 293T cells were transduced with FLAG tagged DDX3X WT, R534H, or 38-44ala. Immunoprecipitation was conducted for FLAG and immunoblotted for ribosome-related proteins and controls. **(C)** Comparison of eIF4E-binding-like motifs. The consensus eIF4E interacting protein motif is compared to the motifs found in 4EBP1 and in DDX3X **(D)** ^15^N HSQC spectrum for m7G bound eIF4E (in red) alone. **(E)** HSQC spectra of m7G bound eIF4E alone (in red) and with the addition of 1mM 4EBP1 peptide (blue) **(F)** HSQC spectra of m7G bound eIF4E alone (in red) and with the addition of 1 mM DDX3X peptide (green). **(G)** Immunofluorescence for DDX3X WT and mutants. Images are representative of N > 10 cells. HEK 293T cells were transduced with DDX3X-FLAG construct, fixed, and stained for anti-DDX3X and the nuclear marker DRAQ7 (Figure 4G).

DDX3X could interact with the translational machinery either through the functional core, one or both termini, or multiple locations. One factor that suggests this interaction is happening in the termini is the possible conservation of this binding between DDX3X and its yeast homolog Ded1^16^. Taken together, these data suggest that the N- and C-termini of DDX3X play a functional role and may potentially mediate interaction with the ribosome.

To determine which regions in DDX3X interact with the translation machinery, we transduced HEK 293T cells with FLAG-tagged full length DDX3X or a truncated 132-607 mutant, and conducted FLAG immunoprecipitation. Full length DDX3X, but not the 132-607 truncation, co-precipitated small ribosomal subunit protein RPS11 and eIF4A, suggesting a loss of interaction with the translation machinery for the truncated mutant (Supplementary Figure 4A).

Having identified the N- and C-termini as locations potentially mediating binding to the ribosome, we then attempted to narrow down the location of the ribosome-binding domain. We made constructs of FLAG-tagged DDX3X with alanine substitutions across termini residues conserved across the DDX3X family, including amino acids 14-21, 38-44, and 600-603 (Supplementary Figure 4B). These constructs were transduced into HEK 293T cells from which RNase-treated lysates were obtained and used for FLAG immunoprecipitation. DDX3X WT pulled down small ribosomal subunit protein RPS11 and translation factors eIF4E and eIF4A, which are associated with the scanning ribosome^35^. Conversely, the different mutants lost interaction with these proteins to different degrees. Both 14-21 and 600-603 mutant showed a similar ability to pull down eIF4E and eIF4A, but a reduced ability to pull down RPS11. DDX3X R534H, a missense mutant found in the functional core which causes helicase activity deficiency^15,19^, pulled down RPS11 to the same degree as wild type, but lost interaction with eIF4E and eIF4A. Finally, the mutant with alanine substitutions spanning the 38-44 residues, which we refer to as 38-44ala, presented the most severe difference from wild type, with greatly reduced copurification of RPS11, eIF4E, and eIF4A (Figure 4B, Supplementary Figure 4B). This suggests that the 38-44 residues of DDX3X are involved in the interaction with the ribosome, whether directly or indirectly through other intermediary proteins.

The 38-44 residues (YIPPHLR) are conserved across the DDX3X/Ded1 subfamily but absent in other DEAD box proteins (Figure 4A)^1^. This sequence has been previously identified as potential eIF4E-binding region because it is reminiscent of the consensus eIF4E-binding sequence YXXXXLΦ found in several eIF4E-binding proteins such as the 4EBPs and eIF4G (Figure 4C). substitution of Tyr38 or Leu43 to alanine partially ablated co-immunoprecipitation of DDX3X and eIF4E in HeLa lysate^36^, in a fashion similar to our observation of the lack of co-precipitation between 38-44ala and eIF4E. We hence decided to test whether these residues mediate DDX3X’s indirect interaction with ribosome via direct binding to eIF4E.

We used NMR spectroscopy to test whether the 38-44 region of DDX3X interacts with eIF4E directly. We purified ^15^N labeled S. cerevisiae eIF4E, which shares the core conserved peptide binding site with human eIF4E^37^. The ^15^N HSQC spectrum of eIF4E bound to m7GTP was well-dispersed, indicating a stable, folded protein (Figure 4D). Addition of 1 mM of a peptide comprising human 4E-BP1 amino acids 51-67 elicited dramatic chemical shift changes (Figure 4E), consistent with the known interaction between 4E-BP1 and eIF4E. In contrast, addition of a peptide comprising human DDX3X amino acids 33-51 caused minimal chemical shift perturbations (Figure 4F). We therefore conclude that human 4E-BP1 binds to S. cerevisiae eIF4E, but that DDX3X amino acids 33-51 do not directly interact with eIF4E.

The lack of co-immunoprecipitation between mutant DDX3X and ribosomal proteins could be due to mutants affecting protein localization. To test this possibility, we used microscopy to determine the localization of the different DDX3X variants. HEK 293T cells were transduced with different DDX3X-FLAG constructs, fixed, and stained for anti-FLAG and the nuclear marker DRAQ7. DDX3X protein was broadly cytoplasmic, and neither the R534H mutation, nor 38-44ala, nor the double mutant 38-44ala/R534H altered this localization compared to WT (Figure 4G). We have therefore identified the 38-44ala residues as contributors to the interaction between DDX3X and the ribosome complex.

### Ribosome-association plays a role in DDX3X function

Having identified a DDX3X mutant that impacts its association with the ribosome, we next investigated the effect of losing this interaction on DDX3X’s translational activity. HCT116 degron cells were transduced with both the HART construct and DDX3X-FLAG-BFP, either WT or helicase defective R534H, 38-44ala, or the double mutant 38-44ala/R534H. FACS was used to obtain a triple-positive population for mCherry, eGFP, and BFP. These cells were treated with auxin for 48 hours and flow cytometry was used to measure the HART ratio.

The HART-ODC1 ratio of auxin-treated cells co-transduced with WT DDX3X was increased compared to parental cells, although not to the same level as control DMSO-treated cells. This suggests the exogenous DDX3X partially rescued the translation defect caused the loss of endogenous DDX3X. Instead, transduction of R534H nor 38-44ala nor the double mutant 38-44ala/R534H not only did not rescue the effect, but further decreased the HART ratio compared to untransduced cells (Figure 5A). Results for HART-RAC1 showed no increase nor decrease in HART ratio in WT, 38-44ala, or R534H/38-44ala cells compared to untransduced, while infection R534H did show a statistically significant decrease compared to untransduced cells (Figure 5B).

**Figure 5:**
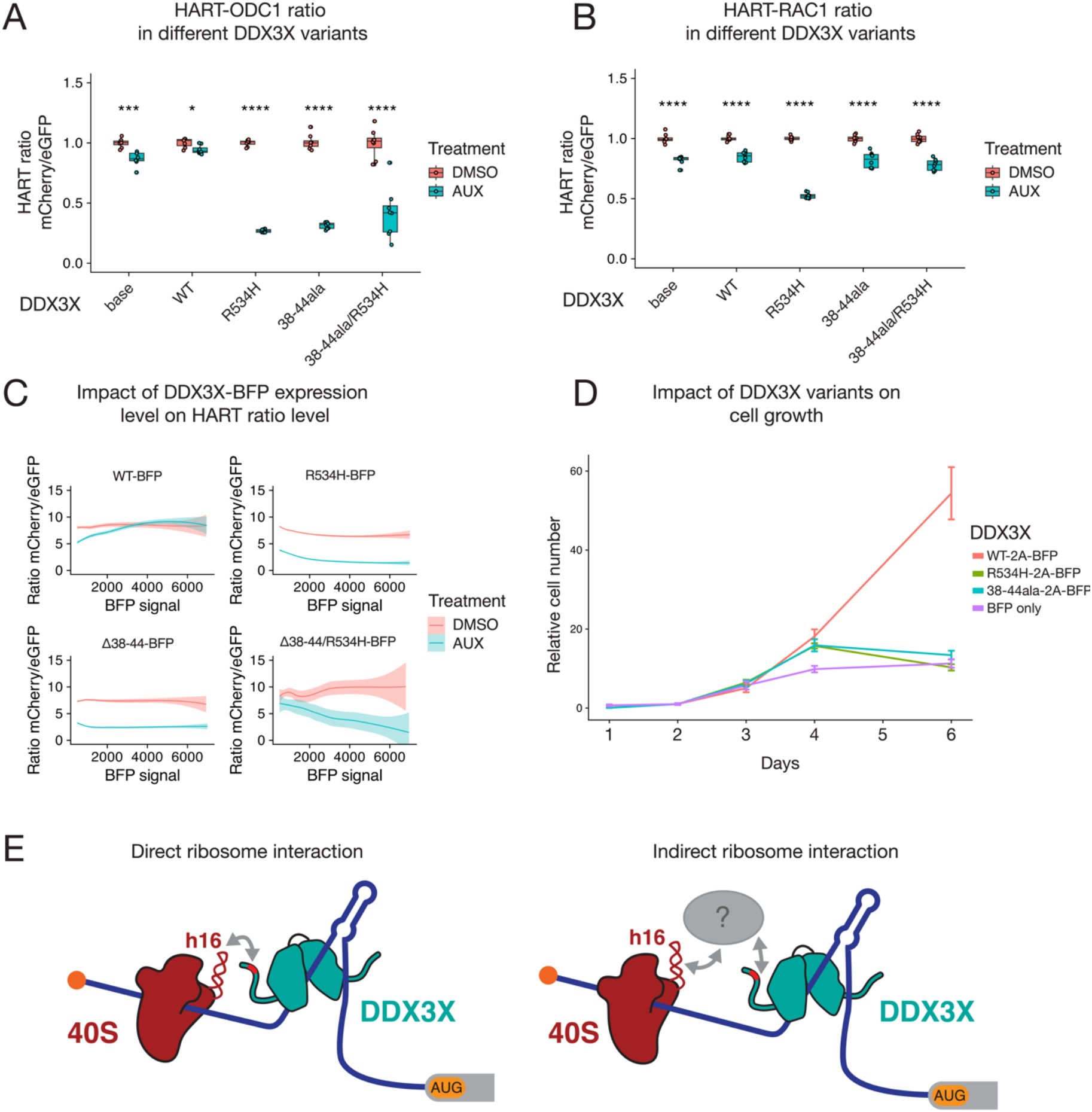
DDX3X 38-44ala cannot rescue translational or cell growth defects caused by the loss of DDX3X. **(A-B)** HART data for indicated DDX3X mutants. HCT116 degron cells were transduced with the HART-ODC1 **(A)** or HART-RAC1 **(B)** construct and different variants of DDX3X-FLAG-BFP, either WT, helicase defective R534H, 38-44ala, or the double mutant R534H/38-44ala. Cells were treated with auxin, which induces loss of the endogenous DDX3X via a degron tag, or DMSO as control. After 48 hours flow cytometry was used to measure fluorescence levels, and the HART ratio (mCherry/eGFP) was calculated. Statistical significance was determined by unpaired *t*-test: ns: p > 0.05, *: p <= 0.05, **: p <= 0.01, ***: p <= 0.001, ****: p <= 0.0001. **(C)** Relationship between DDX3X-FLAG-BFP levels and HART ratio. Flow cytometry was used to measure eGFP and mCherry levels, which constitute the HART ratio, and BFP, which is a proxy for the level of DDX3X-FLAG-BFP expression. **(D)** Cell growth curve for DDX3X variants. HCT116 degron cells were transduced with DDX3X WT, R534H, 38-44ala or BFP control. After auxin was added to degrade endogenous DDX3X, CellTiter Glo was used to measure cell number and plot cell growth. **(E)** Model of DDX3X function in translation and its relation to the ribosome. The 38-44 residues of DDX3X (shown in red) contribute to its interaction with the translational machinery. Helix 16 has been previously identified as mediating this association, but it is unknown whether this interaction happens directly between DDX3X and the ribosome (left) or via intermediary protein(s) such as the components of the eIF4F complex (shown in grey). Our data shows that association between DDX3X and the translational machinery plays a crucial role in DDX3X’s role of unwinding 5′ UTRs to allow ribosome scanning and promote translation of a subset of human mRNAs.

For both reporters, the R534H mutation has a negative effect across 5′ UTRs. This is in line with DDX3X helicase mutations acting as dominant negative in translation assays as well as in human disease^11,12,19,34^. In contrast, the 38-44ala mutant seems to have a dominant negative effect on the ODC1 but a neutral effect on the RAC1 reporter. Interestingly, in both cases the dual mutant R534H/38-44ala followed the pattern of 38-44ala, suggesting that loss of association with the ribosome or associated factors is dominant over helicase mutations in the context of translation. This can be explained in the cis model of DDX3X activity, where interaction with the ribosome is necessary for its role in translation. In the HART-RAC1 data, the helicase defective DDX3X R534H mutant has a dominant negative effect on translation but the 38-44ala mutation rescues this effect. These results suggest that interaction with the ribosome or associated factors plays a role in the translational activity of DDX3X, although different 5′ UTRs are regulated differently.

Lentiviral transduction can lead to variable expression of the exogenous construct across cells. We used the BFP signal, which is a proxy for the expression of DDX3X-FLAG, to investigate the effect of DDX3X expression level on the HART ratio. Notably, higher BFP levels in DDX3X-FLAG-BFP expressing cells correlated with a reduction in the difference of the HART ratio between auxin and DMSO treated cells, indicating dosage sensitivity (Figure 5C). At high levels of BFP, the ratio in auxin-treated cells was above those in DMSO treated cells, suggesting that high enough levels of exogenous DDX3X could promote translation above endogenous levels. This phenomenon was not observed in cells transduced with the mutant DDX3X constructs.

We also investigated the effect of DDX3X mutants on cellular growth. While exogenous DDX3X WT rescued cell-growth defects due to the loss of endogenous DDX3X, neither mutant R534H nor 38-44ala can (Figure 5D). Similarly, the 132-607 truncation of DDX3X, which lacks N- and C-termini but preserves helicase activity^15^, cannot rescue this cell growth phenotype (Supplementary Figure 5A). These data suggest that both helicase function and the association with the ribosome are crucial to DDX3X function in proliferation, although whether this is a result of translational activity or one of the other roles it plays in RNA metabolism is yet to be determined.

Our data shows that the 38-44 residues of DDX3X contribute to its interaction with the translational machinery. This interaction, whether direct with the ribosome or mediated by other factors, is important for DDX3X’s role in unwinding secondary structures in the 5′ UTR of a subset of human mRNAs and promoting their translation (Figure 5E).

## DISCUSSION

In this paper we investigated the translational activity of DDX3X. DDX3X-sensitive 5′ UTRs were selected among transcripts whose ribosome density went down upon loss of DDX3X and whose 5′ UTR was shown to be sensitive to DDX3X^19^. We chose the ODC1 and RAC1 5′ UTRs as they were among the most DDX3X-sensitive transcripts in both in vitro and in cell assays. They also provide diversity in length, at 511 and 197 nt respectively. We used the ODC1 and RAC1 5ʹ UTRs to create a DDX3X translation reporter, termed HART. Notably, HART data replicated the previous assays, with translation of ODC1 and RAC1 5′ UTRs shown to be sensitive to the loss of DDX3X unlike the RPLP1 5′ UTR^19^. Being embedded into the cell as DNA via lentiviral transduction, reporter-encoded mRNAs undergo mRNA processing and export before they are translated. The alignment between HART data and prior in vitro and in cell assays, which are performed with in vitro transcribed and capped mRNA, suggest that the DDX3X-sensitive step is in translation. Differential DDX3X dependence across 5ʹ UTRs attached to the same coding sequence and 3ʹ UTR isolates DDX3X dependence to a step prior to translation elongation.

To promote translation of eGFP in HART and act as an internal control, we selected a short 9 nt 5′ UTR, which is too short to form complex RNA structures that might depend on DDX3X for unwinding before translation. Additionally, the lack of change of eGFP signal between auxin and DMSO conditions in HART (Supplementary Figure 1C) suggests that the loss of DDX3X does not affect its translation. In addition to its lack of complex RNA structure, there could be other reasons that make it insensitive to permutations in DDX3X. Very short 5′ UTRs promote translation in cap dependent and independent mechanisms^38^. Additionally, even in the context of cap dependent translation, they have been observed to direct translation in a ribosome scanning free manner^39,40^. For example, the Translation Initiator of Short 5’ UTR (TISU) is a regulatory element that has a median length of 12 nucleotides and is required for a cap-dependent, scanning-free mechanism of translation initiation on 5′ UTRs with fewer than 30 nucleotides^39^. Notably, TISU function does not require eIF4A, suggesting that some short 5’ UTRs lack the need for DEAD-box helicases^41^. Similarly, cap-assisted translation without scanning was reported for the histone H4 mRNA, which is notable for directing translation in a wide variety of eukaryotic cells^38,42,43^. It is possible that this extremely short 5′ UTR avoids DDX3X dependency by initiating translation without ribosome scanning or by cap independent mechanisms altogether, hence without the need for unwinding of complex structures. Further work into the mechanism of translation of very short 5′ UTR is still needed to understand this phenomenon. Taken together, and in conjunction with HART’s reproducibility in microscopy (Figure 1F), the data suggests that HART is a reliable and versatile reporter for the activity of DDX3X in translation initiation.

One crucial yet incompletely understood aspect of DDX3X regulation is what structures and features make a 5′ UTR translationally DDX3X-sensitive. We used HART to dissect the features of DDX3X-sensitive 5′ UTRs. Both the RAC1 and ODC1 5′ UTRs have high GC content and are predicted by SHAPE-MaP to possess complex secondary structures. This is in line with previous data suggesting high GC content as a marker of DDX3X sensitivity^6,19^ as well data in yeast, where repression of Ded1p activity leads to accumulation of organized RNA structures in 5′ UTRs(Guenther et al., 2018). Elaborate RNA secondary and tertiary structures can sterically block the scanning ribosome, which possesses no intrinsic unwinding activity and relies on helicases such as DDX3X^44^.

Our work suggests that the RAC1 5′ UTR features a large stem loop which relies on DDX3X for unwinding. In the absence of DDX3X, such structure impedes the ribosome’s ability to scan and initiate elongation. Disruption of this structure with deletions of nucleotides 1-39 and 40-78 decreases sensitivity to DDX3X (Figure 3B) and produces 5′ UTRs with lower predicted free energy and structural complexity. Strikingly, the 40-78 deletion causes the reporter to be unaffected by DDX3X loss. Another possibility is that this stem loop regulates uORF usage. Secondary structures in the 5′ UTRs can also increase ribosome dwelling time and promote translation on sub-optimal upstream start codons^45,46^. The RAC1 5′ UTRs features a CUG near-cognate start codon in position 28, right upstream of the stem loop (Figure 3B). In the absence of DDX3X, the stem loop might direct translation towards the uCUG, which might negatively affect translation of the main ORF. This model is consistent with the diminished dependence of DDX3X in the 1-39 and 40-78 deletion constructs. Overall, the data suggests that sensitivity to DDX3X in RAC1 and ODC1 5′ UTRs is conferred by complex secondary structures.

We envision two models of DDX3X unwinding 5′ UTRs structures during ribosome scanning. In the trans model, DDX3X unwinds structures ahead of and untethered from the scanning ribosome and associated translational machinery. In the cis model, DDX3X unwinds the RNA while tethered or bound, similarly to how eIF4A associates with the ribosome during scanning^33^. To differentiate between these two models, we investigated the potential binding of DDX3X to the translational machinery. We showed that full length DDX3X, but not its helicase core alone, can immunoprecipitate the translation machinery (Supplementary Figure 4A), suggesting an interaction between the terminal domains of DDX3X and the ribosome complex. Notably, these termini are highly conserved from yeast to human across the Ded1/DDX3X subfamily, but not in the wider DEAD box family^1,15^. Both DDX3X and Ded1 have been suggested to bind to helix 16 of the 18S of the smaller subunit of the ribosome^16,19^. We reasoned that if this interaction with the ribosome or associated factors is specific to the subfamily, the residues mediating it are likely to be found in the termini.

To determine which residues in DDX3X mediate its interaction with the translation machinery, we made targeted alanine substitutions to several conserved regions in DDX3X termini. Immunoprecipitation showed the 38-44ala mutation, but not DDX3X WT and R534H, lost interaction with ribosomal protein RPS11 and several translation initiation factors (Figure 4B). While it was previously identified as an eIF4E binding region, NMR data showed that the 38-44 residues of DDX3X did not bind directly to eIF4E (Figure 4F). This can be explained by the fact that 38-44 region of DDX3X is similar to the canonical eIF4E binding region found in 4EBP and eIF4G but differs in its last residue, which is a hydrophobic amino acid in the canonical sequence but an arginine in DDX3X(Sachs and Varani, 2000; von der Haar et al., 2004; Richter and Sonenberg, 2005). Residues 38-44 could instead mediate binding to the translational machinery via other factors. Previous work has also suggested that DDX3X interacts with eIF4G(Soto-Rifo et al., 2012; Soto-Rifo et al., 2013; Ayalew et al., 2016; Gong et al., 2021). Direct binding to yeast eIF4G, 4A and 4E was also seen in Ded1(Hilliker et al., 2011; Guenther et al., 2018; Gulay et al., 2020), suggesting interaction between DDX3X and the eIF4F complex(Ryan & Schröder, 2022). We suggest that the 38-44 residues of DDX3X contribute, at least partially, to the interaction with the ribosome, although whether this interaction is direct or mediated by other factors in the translational machinery needs further study (Figure 5E).

In addition to its role in dissecting specific 5′ UTRs sensitivities and DDX3X mutant effects, we anticipate HART to be useful in several other applications. Its reliability as a 5′ UTR-based reporter of function in translation initiation can be employed in conjunction with large naturally occurring or artificial 5′ UTR libraries to gain a granular and large-scale characterization of DDX3X regulation. Alternatively, HART’s readout in both flow cytometry and microscopy setting can be employed to screen for drugs that alter DDX3X behavior or rescue DDX3X or mutation. The lentiviral transduction delivery of HART can be applied to in vivo systems, in other to measure DDX3X function in living systems. Finally, by choosing the proper control and experimental 5’UTRs, it can be adapted to measure the activity of any other helicase or translation factor in addition to DDX3X. HART’s reliability and flexibility can prove a useful tool to study translation in diverse settings, and advance both the understanding of protein production and disease treatment.

## METHODS

### Molecular cloning

DNA amplification was obtained by PCR with either KAPA HiFi HotStart ReadyMix (Roche KK2601) or Q5 High-Fidelity 2X Master Mix (New England Biosystems #M0492) following manufacturer protocol unless otherwise noted. PCR products were digested with 1 μL DpnI (NEB #R0176L) for 1 hour and purified by agarose gel extraction (MinElute Gel Extraction Kit, Qiagen #28606). Plasmid constructs were created by Gibson cloning with NEBuilder HiFi DNA Assembly Master Mix (NEB #E2621L) or Gibson Assembly Master Mix (NEB E2611L). Assembled plasmids were transformed into Mach1 (Invitrogen #C862003) or STBL3 (from Q3 Macrolab at University of California, Berkeley) competent cells for amplification and were purified with QIAprep miniprep (Qiagen #27104).

### DNA sequences

The lentiviral dual promoter plasmid backbone for HART was a gift of the Goodarzi lab and published in Amendola et. al, 2005. The 5′ UTRs of RAC1, HMBS, ODC1, and RPLP1, and control were taken from plasmids from previous work^19^. Deletions in RAC1 and ODC1 were achieved by Gibson cloning. The mutated d4 ornithine decarboxylase (ODC) degron fused to mCherry was inserted via primers with Gibson cloning based on the published sequence: (AGCCATGGCTTCCCGCCGGAGGTGGAGGAGCAGGATGATGGCGCGCTGCCCATGTCTTGTGCCCAGGAGAG CGGGATGGACCGTCACCCTGCAGCCTGTGCTTCTGCTAGGATCAATGTG)^23^. Gibson cloning was used to insert all the HART elements into the lentivirus UCOE-SFFV backbone. The final HART construct is: PuroR-T24-eGFP-[control5′ UTR]-miniCMV-hPGK-[DDX3X-sensitive5′ UTR]-mCherry-d4ODCdegron.

Plasmids were deposited on Addgene:

#207398: KW54 (HART-ODC1)

#207399: KW57 (HART-RAC1)

#207400: KW63 (HART-RAC1-PuroR)

#207401: KW78 (HART-ODC1-PuroR)

Lentiviral DDX3X constructs were created by inserting the DDX3X sequence from previous work^19^ into the UCOE-SFFV backbone (gift from the James K Nunez lab) via Gibson cloning. Lentivirus packaging plasmids included gag/pol, REV, TAT, and VSVG (gift from the Weissman lab). Gibson cloning was also employed to generate the various mutations via primers containing said mutations. BFP and Puromycin resistance sequences were taken from pHR-SFFV-dCas9-BFP-KRAB (Addgene #46911) and Twist respectively. The final DDX3X lentiviral construct sequence is: UCOE-SSFV-DDX3X-3xFLAG-2xP2A-TagBFP-PuromycinR

### Cell culture

HCT116 cells were cultured in McCoy’s 5A media (Gibco #16600082) and 10% FBS (Avantor #97068-085) and 1X Penicillin-Streptomycin Solution (Corning #30002CI). HEK 293T cells were cultured in DMEM with 4.5 g / L glucose, L-glutamine & sodium pyruvate (Corning #10-0130CV) and 10% FBS (Avantor #97068-085) and 1X Penicillin-Streptomycin Solution (Corning #30002CI). Cells were cultured at 37 °C in 5% CO^2^. Cell passage was conducted with trypsin (Corning #25-053-CI). Cells were washed with 1X PBS pH 7.4 (Gibco #70011044) before media change and passaging.

### Lentivirus packaging and infection

For packaging lentivirus, HEK 293T cells were kept at confluency below 90% for several passages. On day 1, HEK 293T cells were plated at 1.2×10^6^ cells/well in 6 well plates. On day 2 cells were transfected with lentiviral DNA following the Mirus-LT1 (Mirus MIR 2300) transfection protocol. Briefly, 6 μL transfection reagent Mirus-LT1 were mixed with 250 μL Opti-MEM (Thermo Fischer #31985062) and incubated at RT for 15 min. 1.5 μg of the desired lentiviral DNA with 0.1 μg gag/pol, REV, and TAT packaging plasmids and 0.2 μg VSVG packaging plasmid (for a total of 2 μg of DNA) were added to the Opti-MEM mix. The mix was incubated at RT for 15 min, and then gently added to the HEK 293T cells dropwise. On day 2, the virus-containing supernatant was harvested by gentle pipetting and filtered with a low protein binding syringe filter (0.22 μM, EDM Millipore # SLGV033RB). The virus was frozen at -80 °C for long-term storage or transduced directly into the desired cells.

Cells to be transduced were plated in 24 well plates at 0.168×10^6^ cells/well on day 1. On day 2, the lentivirus supernatant collected previously was added to the cells. Different amounts of virus, ranging from 5 to 500 μL was added together to Polybrene (EMD Millipore TR-1003-G) to each cell. The plates were spun down for 2 hours at 1000 RPM at 37 °C. On day 4, cells were sorted as described below. The appropriate amount of virus was determined by sorting cells from wells with infection rates below 30%.

Cells in Figures 1 and 3 were transduced with HART constructs containing the indicated DDX3X-sensitive 5′ UTR. Cells in Figure 4 were transduced with DDX3X constructs. Cells in Figure 5 were transduced both with the indicated HART and DDX3X constructs.

### Flow cytometry

For cells transduced with the HART constructs in Figures 1 and 3, FACS was used to sort cells positive for both eGFP and mCherry. For cells transduced with the DDX3X constructs in Figure 4, cells positive for BFP were sorted. For cells transduced with both HART and DDX3X constructs in Figure 5, cells positive for eGFP, mCherry, and BFP were sorted.

FACS was performed using Fortessa SORP to measure the fluorescent signal of eGFP (488nm 50mW laser, Blue B or Blue C), mCherry (561nm. 50mW laser, YG C detector) and BFP (405nm 100mW laser, Violet F detector). Untransduced cells were used as negative controls for gating after filtering for single cells.

Sorted cells were used for the HART experiments in Figures 1, 3, and 5. Cells were plated in 24-well plates at 0.168×10^6^ cells/well with 2 mL of media. 500 mM Auxin (Indole-3-acetic acid) in DMSO (Corning #25950CQC) or DMSO alone were added at a 1:1000 dilution to the media. After 48 hours, cells were trypsinized with 200 μL trypsin and resuspended with 200 μL media. Cells were then flowed with a Fortessa X20 Dual SORP. Untransduced cells were used as negative controls for gating after filtering for single cells. Cells were gated as single cells and double positive for eGFP (488nm 60mW laser, Blue C detector) and mCherry (561nm 50mW laser, YG C detector) for Figures 1 and 3, and in as triple negative for eGFP, mCherry, and BFP 405nm 100mW laser, Violet F detector) in Figure 5. The raw value of the three channels was exported. The ratio of mCherry/eGFP was calculated as the HART readout.

### Cell growth curves

Cells were plated in opaque 96-well plates at 5000 cells per well. Cell number was measured at the indicated time intervals with CellTiter-Glo® Luminescent Cell Viability Assay (Promega #G7571) following manufacturer’s instructions.

### RNA folding prediction

RNA folding predictions were computed using the ViennaRNA Package (Version 2.5.1)^47,48^.

### SHAPE-MaP

SHAPE-MaP was conducted as described^30,31^. First, we in vitro transcribed RNA as described^21^. RAC1 and ODC1 5′ UTRs were placed under control of a T7 promoter and in front of luciferase ORF and transcribed using T7 polymerase^19,21^. For Shape-MaP, 1000 ng of the transcribed RNA in 12 μL water was denatured at 95 °C for 2 min and then snap cooled on ice for 2 min. 6 μL of the 3.3x folding buffer (333 mM HEPES pH 8.0, 333 mM NaCl, 33 mM MgCl2) were added to each sample and the RNA was folded at 37 °C for 20 min. For the unfolded control, 1000 ng RNA was placed in 10 μL formamide, 2 μL 10x DC buffer (500 mM HEPES (pH 8.0), 40 mM EDTA), 6 μL water and denatured at 95 °C for 2 min.

For RNA modification, 1 μL of either DMSO or 2M NAI was added to the RNA and mixed. The folded RNA was incubated 15 min at 37 °C, while the unfolded RNA was incubated 15 min at 60 °C. Added 2 μL 100 mM DTT to all tubes to quench the reaction. Added water to volume of 50 μL and cleaned up the RNA (Zymo RNA clean and concentrator -5 # R1013) and eluted in 15 μL water.

For reverse transcription, 1.5 μL 200 ng/μL of random reverse transcription primers was added to the 15 μL RNA tubes and the reaction was incubated at 65 °C for 10 min and then at 4 °C for 2 min. Fresh MaP buffer was prepared by mixing equal volumes 5x MaP prebuffer (250 mM Tris (pH 8.0), 375 mM KCl, 50 mM DTT, 2.5 mM dNTP each) and 30 mM MnCl_2_. Added 12 μL 2.5x MaP buffer to the RNA tubes and mixed and incubated at 25 °C for 2 min. Added 1 μL Superscript II reverse transcriptase (Thermo Fischer Scientific # 18064014) and incubated at 25 °C for 10 min, then at 42 °C for 90 min, then 10 cycles of 50 °C 2 min and 42 °C 2 min, and then 70 °C for 10 min to deactivate the reverse transcriptase. The reactions were placed on ice and then purified using Zymo DNA clean and concentrator kit -5 (Zymo #D4004) following cDNA protocol. The cDNA was then amplified by PCR and sequenced using Amplicon-EZ (Azenta Lifesciences). ShapeMapper and SuperFold packaged were used to analyze the data, convert it into mutational profiles, create SHAPE reactivity plots, and model RNA secondary structures^49^. VARNA software was used for the automated drawing, visualization and annotation of the secondary structures^50^.

For the in cell SHAPE-MaP, HART-ODC1 and HART-RAC1 cells were grown to ∼80% confluency in 24-well plates. Cells were washed with PBS. Added 150 μL 300 mM NAI in PBS to each well and added the same volume of DMSO in PBS to control cells. The reaction was incubated at 37 °C for 30 min. 200 μL of quench solution (700 mM 2-mercaptoethanol in 1x PBS, made immediately before use) was added to the reaction and incubated for 2 min. Cells were washed with PBS and transferred to into a 1.5 mL tube, then spun down and pelleted and washed again once with PBS. The RNA was extracted using Direct-zol RNA Miniprep Kits (Zymo #R2050) following manufacturer’s instructions (including DNA degradation) and resuspended in 88 μL water. Reverse transcription, library amplification, sequencing, and data analysis were achieved as described above for the in vitro protocol.

### NMR spectroscopy

Isotope labeled His-MBP tagged S. cerevisiae eIF4E was expressed in BL21-star E. coli grown in M9 minimal media with 15N ammonium sulfate as the sole nitrogen source. Induction was performed by addition of 1 mM IPTG when the cultures reached an OD of 0.5 and overnight incubation at 16 °C. Bacteria were lysed by sonication in lysis buffer containing 500 mM NaCl, 0.5% NP-40 (Igepal), 10 mM Imidazole, 20 mM HEPES pH 7.5, 10 mM 2-mercaptoethanol, clarified by centrifugation at 40,000 g, and the clarified lysate was applied to nickel resin (Qiagen) pre-equilibrated in wash buffer (500 mM NaCl, 20 mM Imidazole, 10 mM 2-mercaptoethanol, and 20 mM HEPES pH 7.5). After three washes in wash buffer, purified protein was eluted in elution buffer containing 500 mM NaCl, 250 mM Imidazole, 100 mM Na2SO4, 50 mM sodium phosphate buffer at pH 7.5, and 10 mM 2-mercaptoethanol. Protein was then dialyzed into two liters of buffer containing 20 mM HEPES pH 7.5, 125 mM KCl, and 0.5 mM TCEP overnight at 4 °C while simultaneously the affinity tags were cleaved through addition of 1:40 w/w TEV protease. Dialyzed protein was applied to m7G cap resin (GE Healthcare) equilibrated in dialysis buffer and eluted using dialysis buffer supplemented with 0.1 mM m7GDP^51^. Purified eIF4E protein was exchanged into 50 mM phosphate pH 6.5 and 50 mM KCl using a desalting column^52^, concentrated to 200 μM, and m7GTP was added to 250 μM. DDX3X 33-51 peptide (ASKGRYIPPHLRNREATKGAA) was synthesized by the HHMI Mass Spectrometry Facility at UC Berkeley and dissolved at 50 mM in water, while 4E-BP1 peptide (RIIYDRKFLMECRNSPV; amino acids 51-67) was synthesized by Elim Bio and dissolved at 50 mM in DMSO. Peptides were added to a final concentration of 1 mM. 15N HSQCs were acquired at 298 K using the 900 MHz spectrometer at the Central California 900 MHz NMR facility at UC Berkeley using 10% D2O as a lock signal. Spectra were overlaid using UCSF Sparky.

### Immunoprecipitation

For Figure 4B, HEK 293T cells transduced with DDX3X WT and mutants were washed with cold PBS (Gibco #10010049) and lysed in 500 μL of ice-cold immunoprecipitation buffer (40 mM HEPES–KOH pH 7.5, 100 mM KCl, 1 mM EDTA, 10 mM β-glycerophosphate, 10 mM NaF, 2 mM Na3VO4, 0.4% NP-40 and 1 mM PMSF). The lysates were passaged 5 times through a 25 gauge needle and centrifuged to remove the nuclei. RNAse A (5 μg/mL, Qiagen #19101) was added to the whole cell extracts (WCE) and incubated on ice for 15 min. WCEs were pre-cleared with Mouse IgG-Agarose resin (Sigma-Aldrich #A0919) for 2 hours at 4 °C and then incubated with anti-Flag M2 affinity gel (Millipore Sigma # A2220) for 1.5 hours at 4 °C under continuous rotation. The beads were collected and washed five times with the same immunoprecipitation buffer, and the bead–bound proteins were resolved by SDS-PAGE and analyzed by Western blotting.

For Supplementary Figure 4, HEK 293T cells transduced with DDX3X WT and mutants were washed with cold PBS, collected, and lysed in NP-40 buffer (150 mM NaCL, 1% IGEPAL, 50 mM Tris-Cl p.H. 7.5) with protease inhibitor (Roche #04693132001, cOmplete™ EDTA-free Protease Inhibitor Cocktail). Pipetted up and down, don’t vortex. Samples were placed 45 min at 4 °C and then spun down in centrifuge at 4700 rpm for 2 min. The supernatant was transferred to new tubes and treated with MNAse (ThermoFischer #EN0181) at 1 μL /200 μL lysate for 30 min on the rotor at RT. RNAsin (Fisher Scientific #PRN2615), 4x the amount of MNAse, was used to stop the reaction for 15 min at RT. Anti-FLAG® M2 Magnetic Beads (Sigma-Aldrich #M8823-1ML) were added and used to precipitate FLAG according to protocol (M8823, Millipore). Elution was achieved with 3X FLAG® peptide (F4799, Sigma-Aldrich,). Briefly, 30 μL of FLAG peptide (150 μL/mL in TBS with protease inhibitor) was added to the tube and placed on rotor for 30 min. The supernatant was collected and resolved by SDS-PAGE and analyzed by Western blotting.

### Western blotting

Primary antibodies used in this study include rabbit polyclonal anti-DDX3X (custom made by Genemed Synthesis using peptide ENALGLDQQFAGLDLNSSDNQS; Figure 4), anti-actin HRP (Santa Cruz Biotechnology, sc-47778), anti-FLAG HRP (Sigma, A8592), anti-eIF4A1 antibody (ab31217), anti-eIF4G1 antibody (Cell signaling technologies #2498), anti-eIF4E antibody (Cell signaling technologies #9742), anti-ribosomal protein S11/RPS11 (ab175213), anti-RPL10A (Abcam #ab174318).

### Microscopy

For microscopy in Figure 1, HCT116 degron cells transduced with HART-ODC1 or HART-RAC1 were plated in a 384 well plate (Perkin Elmer #6057302) and were treated with AUX or DMSO for 96 hours. Media was removed and 50 μL of 4% PFA in PBS was added to each well for 15 min and incubated at RT. PFA solution was removed and the cells were washed with 100 μL PBS three times using a BioTek EL406. Cells were incubated with 50 μL 1X DAPI (Biotium #40043) in PBS for 5 min at RT. Cells were washed with 100 μL PBS three times using a BioTek EL406 and imaged using a InCell Analyzer 6500HS (GE), taking three fields-of-view per well. HART ratio was calculated as the ratio between the mCherry and GFP channels.

For microscopy in Figure 4, HEK 293T cells were grown in 96 well plates. When they reached 70% confluency media was removed and cells were washed once with PBS. 50 μL of 4% PFA in PBS was added to the well for 10 min at RT. Cells were washed three times with ice-cold PBS and then incubated for 10 min in PBS containing 0.1% Triton X-100 at 4 °C by rocking. Cells were washed in PBS three times for 5 min at 4 °C by rocking and then incubated with 1% BSA PBST solution for 30 min at 4 °C by rocking. Cells were incubated in diluted antibody (ANTI-FLAG, 1:1000, Sigma Aldrich #F1804) in 1% BSA PBST overnight at 4 °C by rocking. Cells were washed in PBS three times for 5 min at 4 °C by rocking and then incubated by rocking with secondary antibody (Goat-anti-Mouse 488, 1:2000, Invitrogen #A-11008) in 1% BSA PBST for 1 hour at RT in the dark. Cells were washed in PBS three times for 5 min at 4 °C by rocking in the dark. 1X DRAQ7 (Thermo Fisher Scientific #D15106) in PBS was added to the cells for 15 min by rocking, followed by one wash with PBS. Images were taken with a confocal microscope at a 20X-40X magnification.

## ACKNOWLEDGMENTS

We thank all members of the Floor Lab for their invaluable support, expertise, and feedback. We are indebted to the UCSF Parnassus Flow Cytometry CoLab staff for their technical assistance and expertise (RRID:SCR_018206). Research reported here was supported in part by the DRC Center Grant NIH P30 DK063720 and by the NIH S10 Instrumentation Grant S10 1S10OD021822-01. This work was supported by the DDX3X Foundation (to S.N.F.) and the National Institutes of Health R35GM149255 (to S.N.F.) and R01NS120667 (to S.N.F.) and the California Tobacco-Related Disease Research Grants Program 27KT-0003 (to SNF) and T30DT1004 (to KCW). S.N.F. is a Pew Scholar in the Biomedical Sciences, supported by The Pew Charitable Trusts. We thank David King for help with peptide synthesis and general advice.

## AUTHOR CONTRIBUTIONS

Conceptualization, K.C.W. and S.N.F.; Investigation, K.C.W., T.S., S.G., J.R.D., S.N.F. Writing – K.C.W. and S.N.F.; Funding Acquisition, K.C.W., S.N.F.; Supervision, S.N.F.

---

**Supplementary Figure 1.**
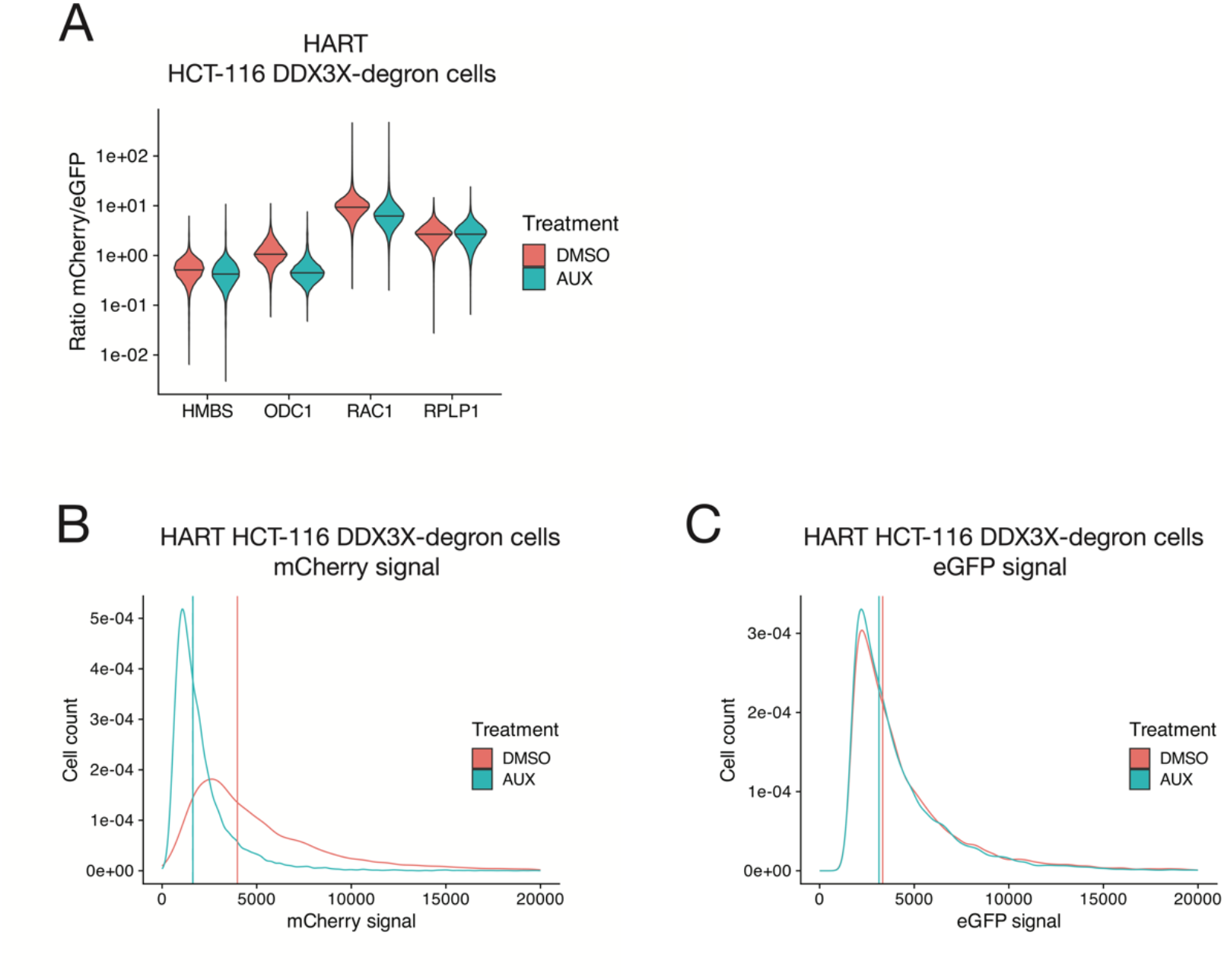
**(A)** Violin plot for HART ratio in HCT116 degron cells analyzed with flow cytometry for the experiment in Figure 1B. HCT116 degron cells were lentivirally transduced with HART constructs with various 5′ UTRs in front of mCherry. After 48 hours from the addition of either DMSO or auxin, which induces degradation of endogenous DDX3X, the fluorescent signal was measured by fluorescent cytometry. The HART ratio (mCherry/eGFP) was calculated for each cell and plotted as a violin plot. **(B)** Flow cytometry data for the mCherry channel for experiment in Figure 1B. The raw data for the 561nm 50mW laser, YG C detector (corresponding to mCherry) for each cell was plotted and the mean was calculated and plotted as a vertical line. **(C)** Flow cytometry data for the eGFP channel for cells in Figure 1B. The raw data for the 488nm 60mW laser, Blue C detector (corresponding to eGFP) for each cell was plotted and the mean was calculated and plotted as a vertical line.

**Supplementary Figure 2.**
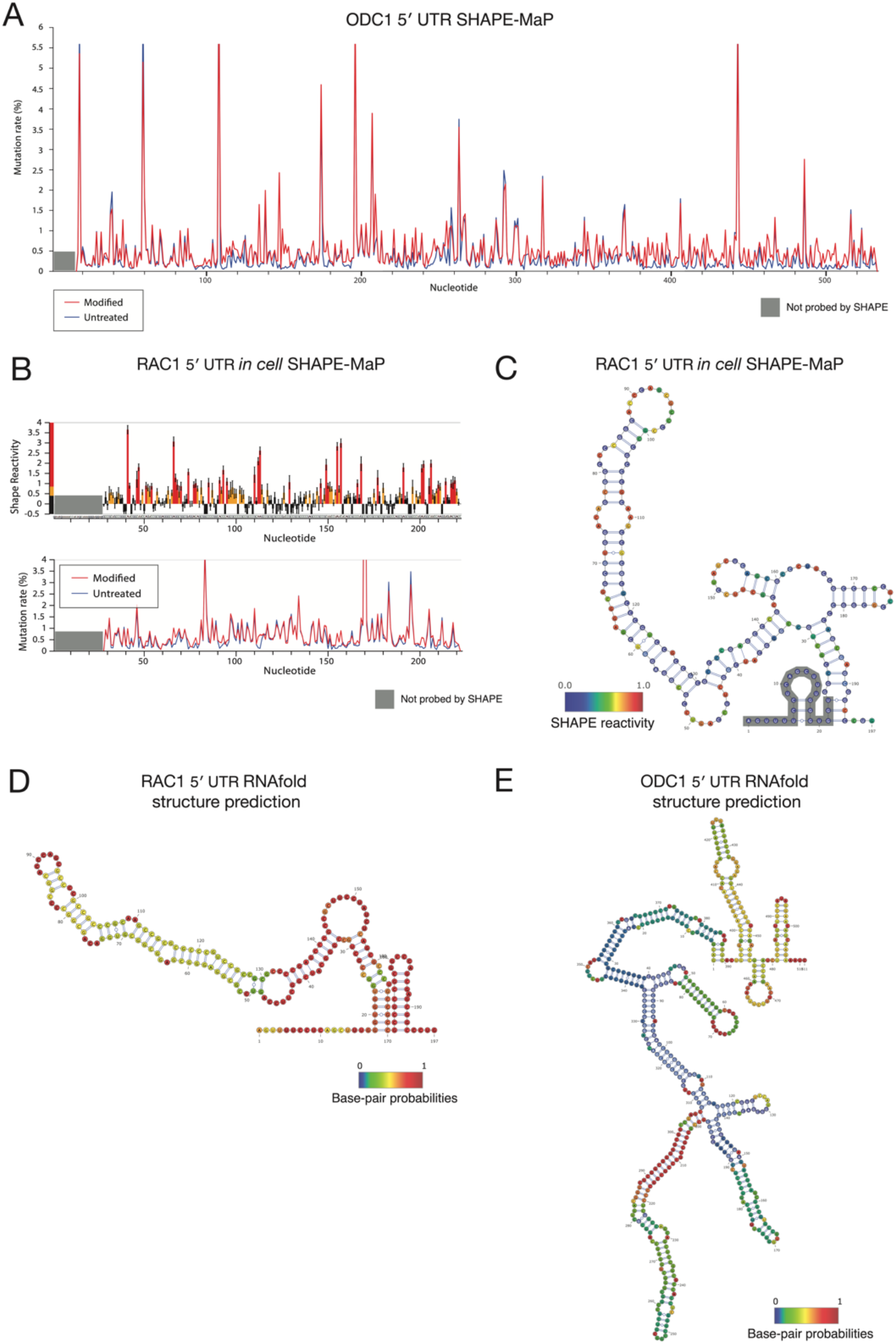
**(A)** SHAPE-MaP reactivity for the 5′ UTR of RAC1 *in vitro* from Figure 2C. *In vitro* transcribed mRNA containing the 5′ UTR of ODC1 and the open reading frame of luciferase was probed with 200 mM NAI or DMSO control for SHAPE-MaP. The RNA was reverse transcribed and sequenced. The SHAPE reactivity was calculated based on the difference in mutation rate. **(B)** SHAPE-MaP reactivity and mutation rate for the 5′ UTR of RAC1 *in cell*. Cells were treated with 300 mM NAI in PBS or control for 20 min before quenching the reaction and extracting the RNA. The RNA was reverse transcribed, sequenced, and analyzed to obtain mutation profiles and SHAPE reactivity with the ShapeMapper tool. **(C)** Diagram of the structure of the RAC1 5′ UTR *in cell*, based on data from Supplementary Figure 2A and computed with ShapeMapper 2.1.3. (Busan & Weeks, 2018) **(D-E)** RNA folding minimum free energy prediction of the structures of the 5′ UTRs of RAC1 (C) and ODC1 (D) using the ViennaRNA Package (Lorenz et al., 2011). The minimum free energy for the RAC1 structure is -97.20 kcal/mol and for the ODC1 is -251.70 kcal/mol.

**Supplementary Figure 3.**
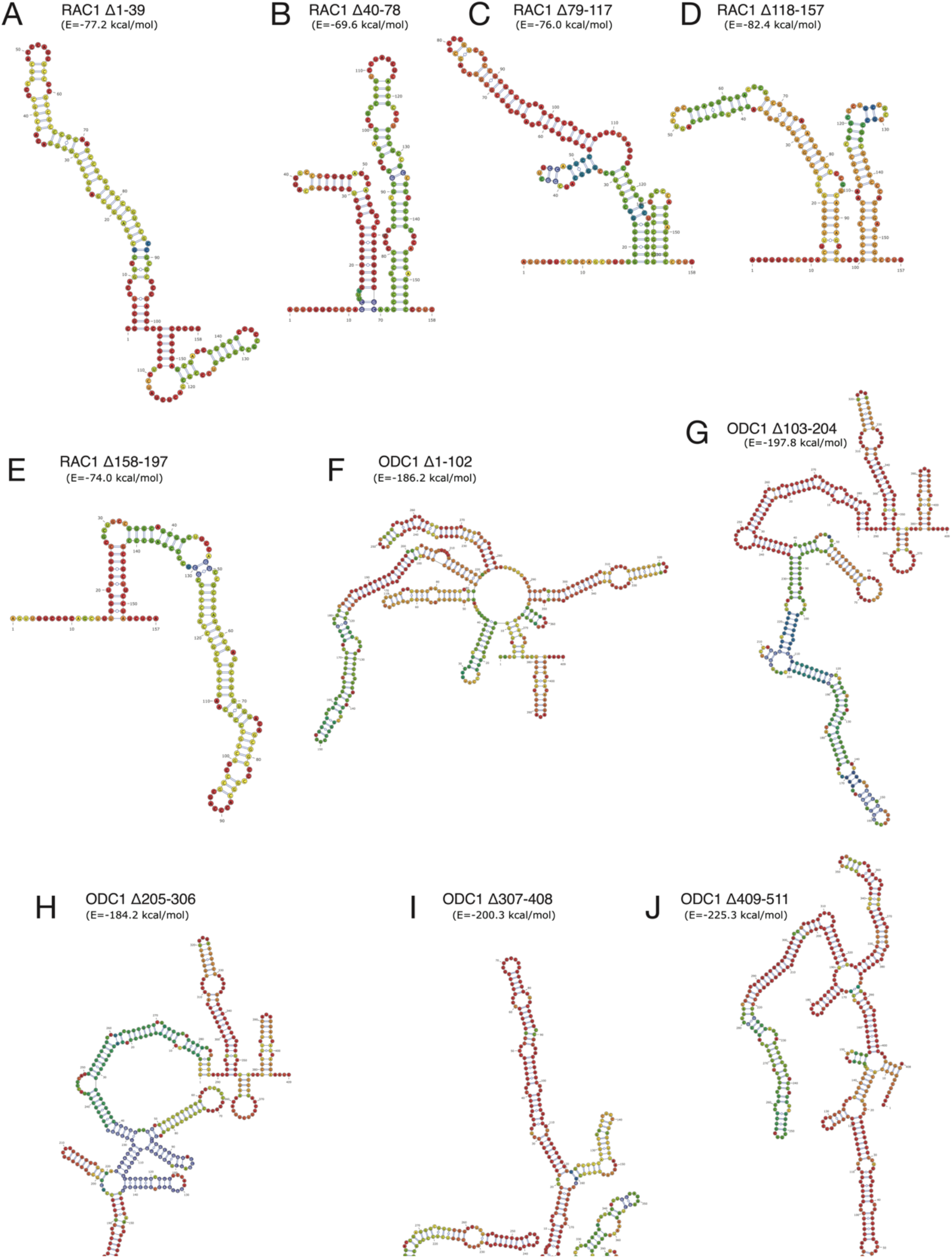
**(A-E)** RNA folding minimum free energy prediction of the structures of the 5′ UTR of RAC1 containing the deletions used in Figure 3 B-C using the ViennaRNA Package (Lorenz et al., 2011). The predicted minimum free energies for the RAC1 deletion constructs are: -77.20 kcal/mol for Δ1-39 -77.20 kcal/mol, -69.60 kcal/mol for Δ40-78, -76.00 kcal/mol for Δ79-117, -82.40 kcal/mol for Δ118-157, -74.00 kcal/mol for Δ158-197. **(F-J)** RNA folding minimum free energy prediction of the structures of the 5′ UTR of ODC1 containing the deletions used in Figure 3 B-C using the ViennaRNA Package (Lorenz et al., 2011). The predicted minimum free energies for the ODC1 deletion constructs are: -186.20 kcal/mol for Δ1-102, - 197.80 kcal/mol for Δ103-204, -184.20 kcal/mol for Δ205-306, -200.30 kcal/mol for Δ307-408, -225.30 kcal/mol for Δ409-511.

**Supplementary Figure 4.**
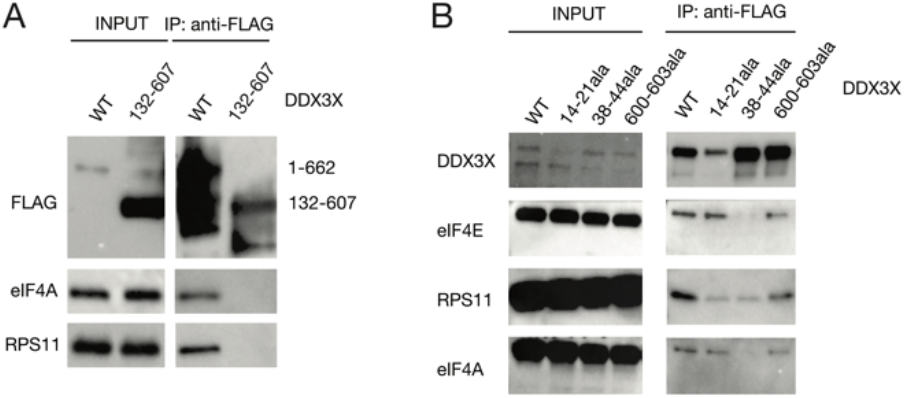
**(A)** DDX3X full length and 132-607 immunoprecipitation. HEK 293T cells were lentivirally transduced with FLAG tagged DDX3X full length or DDX3X 132-607, which represents its functional helicase core. Immunoprecipitation was conducted for FLAG and run on western blot, staining for ribosome-related proteins and controls. **(B)** Immunoprecipitation of DDX3X mutants. HEK 293T cells were lentivirally transduced with FLAG tagged DDX3X WT or several mutants, including the helicase defective mutant R534H and three N- and C-termini mutations in sites conserved across the DDX3X/Ded1 subfamily. Immunoprecipitation was conducted for FLAG and run on western blot, staining for ribosome-related proteins and controls.

**Supplementary Figure 5.**
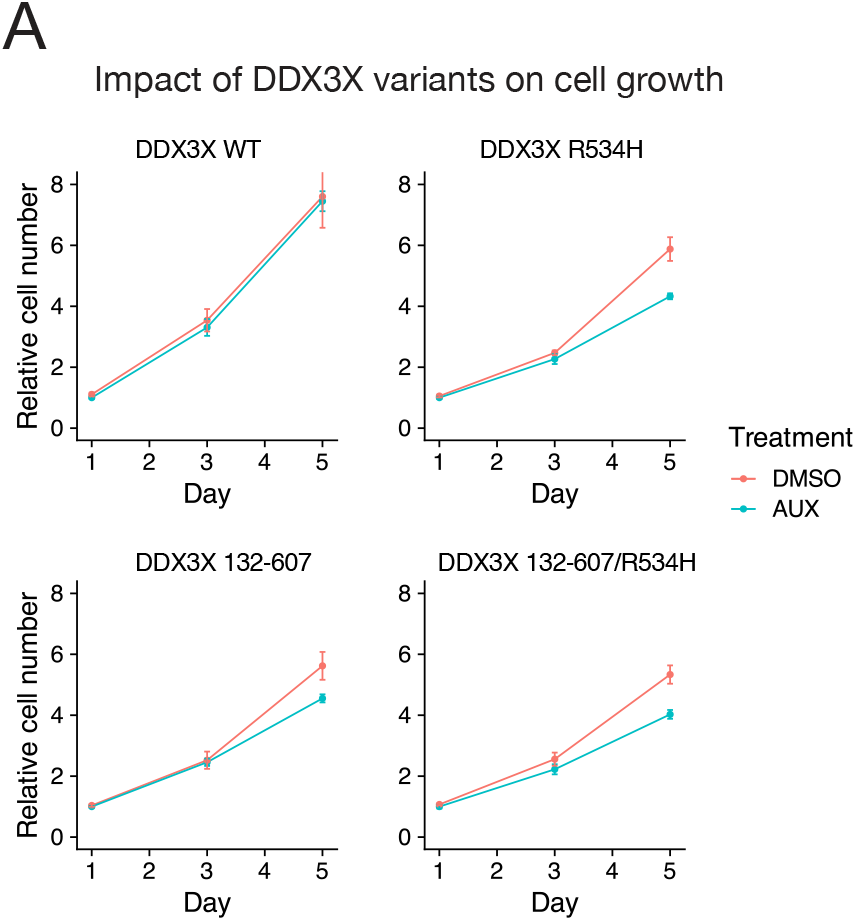
**(A)** Cell growth curves for DDX3X variants. HCT116 degron cells where lentivirally transduced with exogenous DDX3X WT and mutants. Auxin was added to induce loss of endogenous DDX3X and cell number was measured over time with CellTiter Glo.

